# Inflammation and Cellular Stress Induced Neurological Sequelae of *Plasmodium falciparum* Malaria

**DOI:** 10.1101/2021.06.16.448682

**Authors:** Akua A. Karikari, Wasco Wruck, James Adjaye

## Abstract

**Background:** Malaria caused by *Plasmodium falciparum* results in severe complications including cerebral malaria (CM) especially in children. While the majority of *falciparum* malaria survivors make a full recovery, there are reports of some patients ending up with neurological sequelae.

**Methods:** We performed an analysis of pooled transcriptome data of whole blood samples derived from two studies involving various *Plasmodium falciparum* infections, comprising mild malaria (MM), non-cerebral severe malaria (NCM) and CM. Pathways and gene ontologies (GOs) elevated in the distinct *falciparum* infections were identified.

**Results:** Contrary to other research findings, our analysis showed MM share similar biological processes with cancer and neurodegenerative diseases, NCM is associated with drug resistance and glutathione metabolism and CM is correlated with endocannabinoid signaling and non-alcoholic fatty liver disease (NAFLD). GO revealed the terms biogenesis, DNA damage response and IL-10 production in MM, down-regulation of cytoskeletal organization and amyloid-beta clearance in NCM and aberrant signaling, neutrophil degranulation and gene repression in CM. Differential gene expression analysis between CM and NCM showed the up-regulation of neutrophil activation and response to herbicides while regulation of axon diameter was down-regulated in CM.

**Conclusions:** The results of this study have demonstrated that the deleterious effect of *falciparum* malaria on the brain may not be limited to CM and NCM alone but also MM. However, the severity of neurological deficit in CM might be due to the down-regulation of various genes involved in cellular function through transcriptional repression, axonal dysfunction, dysregulation of signaling pathways and neurodegeneration as a result of inflammation and cellular stress. We anticipate that our data might form the basis for future hypothesis-driven malaria research.

## Background

Malaria is a major infectious disease and public health problem with an estimated 215 million cases and 384 000 deaths in 2019 [1]. The disease is most severe in Africa, where the world health organization (WHO) malaria report in 2020 revealed that the region accounts for 94% of malaria cases and deaths universally [1]. Of the five human parasite species, *Plasmodium falciparum* and *Plasmodium vivax* are the most common cause of severe forms of malaria [2]. P. *falciparum* is however predominantly implicated in cerebral malaria (CM) in children under 5 years of age and pregnant women [3–5]. It is also resistant to many anti-malarial drugs making it more lethal.

The life cycle of the apicomplexan *falciparum* within the human host includes the intra-erythrocytic stage where heme is produced from the host’s hemoglobin [6]. *P. falciparum* also harbors the “plant-like” apicoplast which is essential for parasite *de novo* synthesis of heme as a back-up [7, 8]. Heme is used by the parasite mitochondrial electron transport chain and its toxicity is attenuated by crosslinking into an insoluble polymer (hemozoin), via the action of a parasite-specific biochemical activity[6, 9, 10]. Hemozoin (HMZ), also classified as malaria pigment, is typically observed in the liver, spleen and brain of infected individuals [11, 12]. The severity of *falciparum* malaria correlates with the release of HMZ, which can be incorporated into membranes and also diffuse through the blood brain barrier (BBB) [3, 13–15]. Though *P. falciparum* does not invade the brain parenchyma, toxic HMZ crosses the BBB, causing injuries to neurons [16, 17]. Germane to HMZ pathology is the production of reactive oxygen species (ROS), which is implicated in oxidative macromolecular damage, inflammatory response, endoplasmic reticulum (ER) stress and apoptosis [18–22]. The developing brain is especially susceptible to genomic instability and cellular stress engendered by oxidative modification of macromolecules and this is also linked to neurodevelopmental and neurodegenerative diseases including Schizophrenia, Alzheimer’s (AD) and Parkinson’s (PD) diseases [23–27]. Interestingly, reports by Thiam et al. and Cabantous et al. demonstrated the activation of genes associated with AD and PD in CM [28, 29]. In addition, Boldt et al. found a signature of 22 differentially expressed genes related to immunopathological processes and complement regulation in the transcriptome of distinct conditions of childhood malaria including CM [30].

In this study, analysis of two pooled series of microarray of transcriptome data of whole blood cells derived from patients with mild malaria (MM), non-cerebral severe malaria (NCM) and CM were carried out. Our results showed fewer genes expressed in CM compared to the other states of malaria. Additionally, genes involved in neutrophil degranulation and response to herbicides were up-regulated in CM whereas genes associated with axon diameter were down-regulated. Genes connected to DNA repair mechanisms and cytoskeletal destabilization were significantly expressed in MM and NCM respectively.

Thus, the neurological sequelae such as cognitive decline reported in *falciparum* patients, especially CM, could be attributed to inflammation, genomic instability and cellular stress-mediated neuronal malfunctioning and degeneration.

## Methods

### Preprocessing of gene expression data for the pooled analysis

Datasets of the accession numbers GSE1124 and GSE116306 containing transcriptome microarray data of cerebral malaria, non-cerebral severe malaria and mild malaria were downloaded from NCBI GEO. For GSE1124, children within the ages of 0.5 - 6 years were used whereas patients within ages of 1 - 72 years were employed in GSE116306. The number of subjects involved in the study were 4 - 6 (MM:6, NCM:4 and CM:6) for Thiam et al. and 20 samples per each group for Boldt et al. A complete blood count was conducted in both studies and Boldt et al. report that the experimental groups differed significantly with respect to age, respiration rate, degree of spleen enlargement, white blood cell count, glycemia, hemoglobin and hematocrit levels. Thiam et al. on the other hand stated that with the exception of platelet count which differed between the experimental groups, there was no significant difference between the groups for age, hemoglobin concentration, red blood cell count and leucocyte count. The datasets employed in this study are listed in Table 1. GSE1124 was associated with a publication by Boldt et al. [30] and GSE116306 was associated with Thiam et al. [28]. For comparison of distinct disease states including CM, Boldt et al. had processed their data with proprietary software followed by application of Significance Analysis of Microarrays (SAM) on the logarithmic (base 2) signal ratios to identify significantly changed genes at a false discovery rate of 0.004%. We only used the datasets from the accession GSE1124 which were generated on the *Affymetrix Human Genome U133A Array* platform. Datasets from the accession GSE116306 were generated on the *Agilent-039494 SurePrint G3 Human GE v2 8x60K Microarray* platform. All datasets were imported into the R/Bioconductor environment [31].

**Table 1:** Datasets employed in this pooled analysis.

| GEO_sample | Sample name | Group | GEO_accession |
| --- | --- | --- | --- |
| GSM3227838 | CM1 | CM | GSE116306 |
| GSM3227839 | CM2 | CM | GSE116306 |
| GSM3227840 | CM3 | CM | GSE116306 |
| GSM3227841 | CM4 | CM | GSE116306 |
| GSM3227842 | CM5 | CM | GSE116306 |
| GSM3227843 | CM6 | CM | GSE116306 |
| GSM3227844 | MM1 | MM | GSE116306 |
| GSM3227845 | MM2 | MM | GSE116306 |
| GSM3227846 | MM3 | MM | GSE116306 |
| GSM3227847 | MM4 | MM | GSE116306 |
| GSM3227848 | MM5 | MM | GSE116306 |
| GSM3227849 | MM6 | MM | GSE116306 |
| GSM3227850 | NCM1 | NCM | GSE116306 |
| GSM3227851 | NCM2 | NCM | GSE116306 |
| GSM3227852 | NCM3 | NCM | GSE116306 |
| GSM3227853 | NCM4 | NCM | GSE116306 |
| GSM711624 | Asymptomatic_A | Asymptomatic | GSE1124 |
| GSM711625 | Asymptomatic_B | Asymptomatic | GSE1124 |
| GSM711626 | Asymptomatic_C | Asymptomatic | GSE1124 |
| GSM711627 | Asymptomatic_F | Asymptomatic | GSE1124 |
| GSM711628 | Asymptomatic_G | Asymptomatic | GSE1124 |
| GSM711629 | uncomplicated_malaria_D | MM | GSE1124 |
| GSM711630 | uncomplicated_malaria_E | MM | GSE1124 |
| GSM711631 | uncomplicated_malaria_F | MM | GSE1124 |
| GSM711632 | uncomplicated_malaria_G | MM | GSE1124 |
| GSM711633 | uncomplicated_malaria_H | MM | GSE1124 |
| GSM711634 | severe_malaria_A | NCM | GSE1124 |
| GSM711635 | severe_malaria_B | NCM | GSE1124 |
| GSM711636 | severe_malaria_C | NCM | GSE1124 |
| GSM711637 | severe_malaria_D | NCM | GSE1124 |
| GSM711638 | severe_malaria_E | NCM | GSE1124 |
| GSM711639 | cerebral_malaria_C | CM | GSE1124 |
| GSM711640 | cerebral_malaria_D | CM | GSE1124 |
| GSM711641 | cerebral_malaria_E | CM | GSE1124 |
| GSM711642 | cerebral_malaria_F | CM | GSE1124 |
| GSM711643 | cerebral_malaria_G | CM | GSE1124 |

MM: mild malaria, NCM: severe non-cerebral malaria, CM: cerebral malaria. Datasets from the accession GSE1124
are associated with Boldt et al., datasets from the accession GSE116306 are associated with Thiam et al.

The datasets generated on the Agilent platform were read and preprocessed employing the Bioconductor package limma [32]. This included background correction with the method normexp and quantile normalization. The binary score “gIsWellAboveBG” from the input data files was used as binary measure for gene expression. Multiple probes matching the same gene symbol were reduced to the probe with the maximal mean expression signal.

Datasets generated on the Affymetrix platform were read and pre-processed employing the Bioconductor package affy [33]. This included background correction and normalization with the Robust Multi-Array average (RMA) method. Background detection values were calculated via the *mas5calls* method[33]. Analogously to the other platform multiple probes matching the same gene symbol were reduced to the probe with the maximal mean expression signal.

### Analysis of pooled gene expression data

The preprocessed datasets from the GEO accessions GSE1124 and GSE116306 were adjusted for batch effects via the ComBat method [34] from the Bioconductor package sva [35]. In the follow-up analyses this batch-effect-adjusted data was used for differential expression and cluster analysis while the binary expression score described above was used for the Venn diagram analysis. The global cluster dendrogram was generated via the R method *hclust* from the log2-transformed expression data with a coefficient of variation above 0.1 employing Pearson correlation as similarity measure and complete linkage as agglomeration method. Differentially expressed genes were determined with the criteria ratio > 2 and p-value < 0.05 and q-value < 0.25 for up- and ratio < 0.5 and p-value < 0.05 and q-value < 0.25 for the down-regulated genes. The p-value was calculated with the method from the Bioconductor package limma [32]. The p-value was adjusted for multiple-testing with the package qvalue [36]. The heatmaps were generated with the method heatmap.2 from the package gplots [37]. For the Venn diagram analysis genes were considered expressed when the mean of the binary expression score (1 if expressed, else 0) was above a threshold of 0.5 in the Agilent data and the mean of the Affymetrix detection-p-values was below 0.05 in the distinct disease states. The Venn diagrams were generated with the package VennDiagram [38].

### Over-representation analysis of pathways and gene ontologies (GOs)

Pathways annotated with gene symbols were downloaded from the Kyoto Encyclopedia of Genes and Genomes (KEGG) database [39] in July 2020. Genes found in subsets of the Venn diagram analysis and differential gene expression analysis were tested for over-representation in these gene sets via the hypergeometric distribution test implemented in R. Furthermore, these subsets of genes were analysed for over-representation in GOs with the R package GOstats [40]. Dot plots of pathways or GOs most significantly elevated were generated via the R package ggplot2 [41].

### Metascape analysis of differentially expressed genes

Lists of genes differentially up-(ratio > 2, limma-p-value < 0.05, q-value < 0.25) and down-regulated (ratio < 0.5, limma-p-value < 0.05, q-value < 0.25) between CM and NCM were joined to a two-column-list which was subjected to Metscape multi-column-list analysis [42]. From the results the Protein-protein-interaction (PPI) network and the heatmap of regulating transcription factors generated via the TRRUST method [43] were used for this publication.

## Results

We present the outcomes of the pooled microarray datasets of whole blood cells derived from children and adult patients suffering from MM, NCM and CM. Data of gene expression and pathways are represented as dendrograms, Venn diagrams and heatmaps.

### Cluster analysis of CM in comparison to MM and NCM

Fig. 1 shows a categorized cluster of the datasets used in the pooled analysis namely, GSE116306 and GSE1124. In GSE116306, MM corresponds to “mild malaria”, NCM is “severe non-cerebral malaria” and CM refers to “cerebral malaria”. For GSE1124, MM stands for “uncomplicated malaria”, NCM refers to “severe malaria” and CM is “cerebral malaria”. The cluster dendrogram indicates that while there is a general overlap between the different states of *falciparum* malaria, CM cases predominantly share characteristics with NCM when compared to MM.

**Figure 1.**
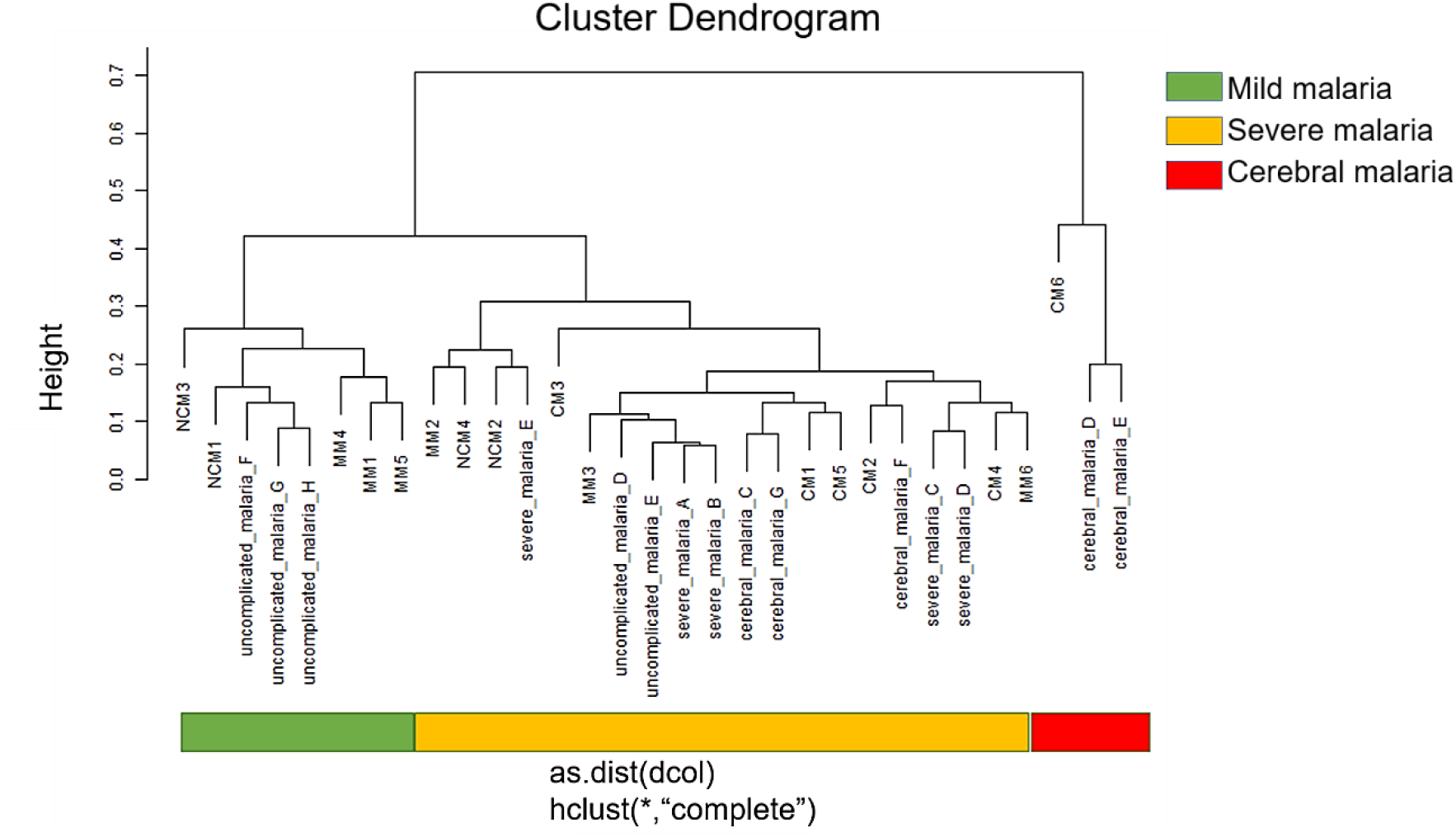
Cluster analysis dendrogram of malaria patient’s transcriptomes displays three clusters with varying severity. The cluster to the right is consistently CM, however contains only three samples, the cluster to the left is more mild, the one in the middle more severe. The distinct clusters are marked with color bars: cluster 1 (green, predominantly mild), cluster 2 (orange, predominantly severe and cerebral), cluster 3 (red, consistently CM). CM (cerebral malaria), MM (mild malaria) and NCM (severe non-cerebral malaria).

### Fewer genes are expressed in CM

Gene expression was analyzed between MM, NCM and CM and represented in a Venn diagram in Fig. 2c. In all, there were 2876 genes mutually expressed between the three disease states. There were 352 and 247 genes independently expressed in MM and NCM respectively. In CM (27 genes) however, several genes were down-regulated which included those associated with cytoskeletal organization, translation, cellular transport, cellular respiration, mitochondrial functioning and proteasomal processes. Genes expressed in common between the 3 classes of malaria comprised those of inflammatory response (CKLF), ER stress (RTNF13), detoxification of toxic metals and antioxidant response (MT1X), cellular stress (HSPs), clathrin and adaptor protein complex 2 assembly (PICALM) and mental retardation (AUTS2). It is noteworthy that genes implicated in AD, (PSEN1 and APP), were also up-regulated in MM, NCM and CM.

**Figure 2.**
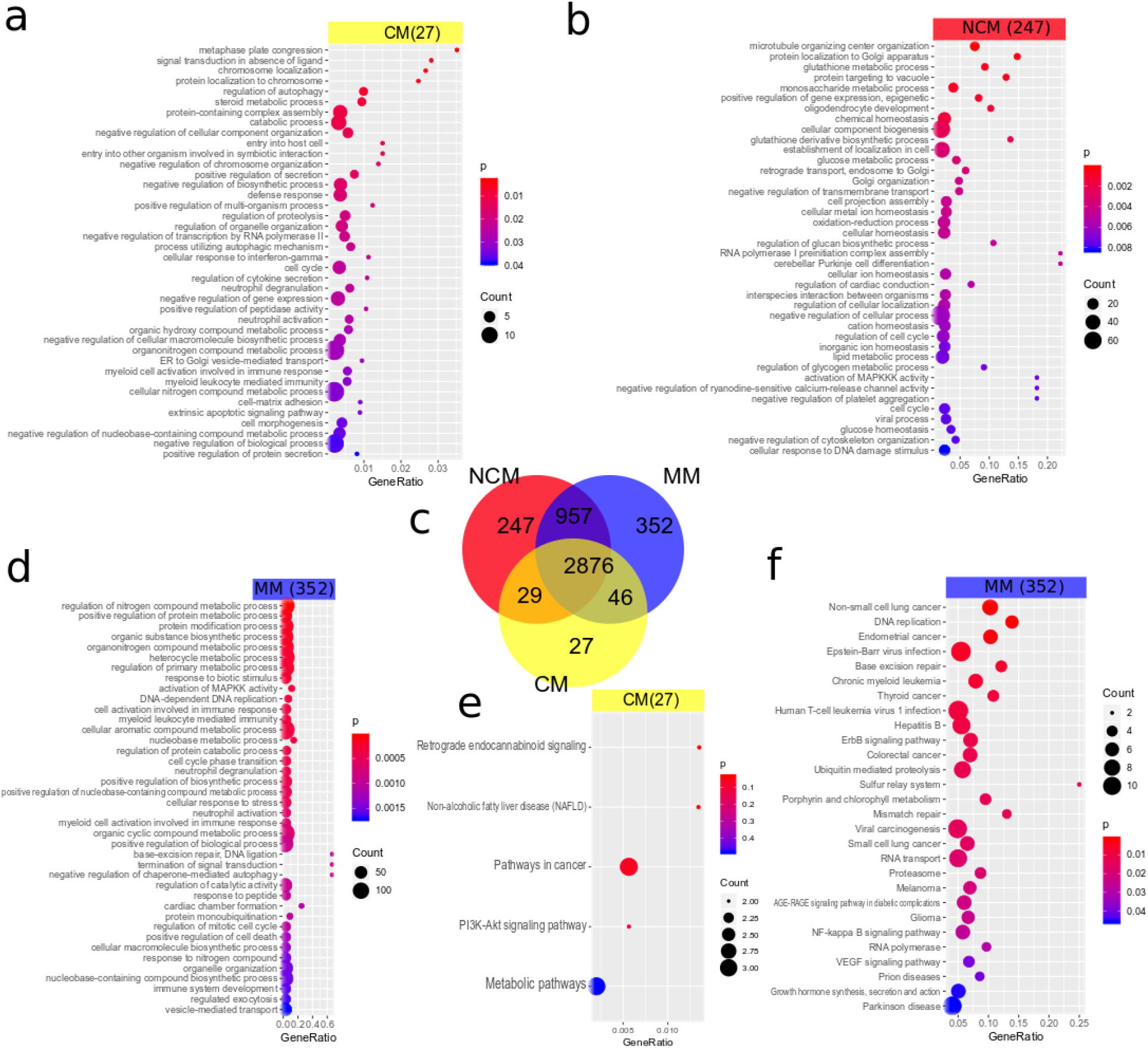
Comparison of gene expression between CM, NCM and MM shows endocannabinoid signaling in CM, neurodegenerative pathways with MM and drug resistance pathways were associated with NCM. Panels (a), (b) and (d) show GO expression analysis results and (e) and (f) show pathway analysis results of the genes expressed exclusively in CM, NCM or MM in the Venn diagram in (c). 27 genes are expressed exclusively in CM while in the NCM there are 247 genes and in MM 352 genes exclusively expressed. Most genes (2876) are expressed in common in all disease states. (a) The GO analysis reveals signal transduction abnormality, neutrophil degranulation and negative regulation of gene expression amongst the most significantly expressed GO terms in the 27 CM genes. (b) Amongst the significant GO terms for the 247 genes expressed exclusively in NCM were *microtubule organizing center organization, glutathione metabolic process, glutathione derivative biosynthetic process, oxidation-reduction process, DNA damage checkpoint, mitotic cell cycle arrest, reactive oxygen species biosynthetic process, removal of superoxide radicals, detoxification, activation of mitogen-activated protein* kinases *(MAP3Ks) activity, negative regulation of cytoskeleton organization and amyloid-beta clearance.* These terms are suggestive of glutathione involvement in defense against ROS in severe malaria, DNA damage repair mechanisms, cytoskeletal destabilization activation and protein clearance. (d) Elevated GO terms for the 352 genes exclusively expressed in MM included *biogenesis*, *regulation of nitrogen compound metabolic process, positive regulation of protein metabolic process, response to biotic stimulus, DNA damage response, detection of DNA damage, positive regulation of phospholipid metabolic process*, *blood vessel remodeling, cell cycle arrest, response to unfolded protein, immune response* and *interleukin (IL)-10 production*, which are indicative of DNA repair mechanisms, cellular repair and anti-inflammatory mechanisms. (e) Over-represented KEGG pathways for the CM-exclusive genes included *retrograde endocannabinoid signaling* and (f) over-represented KEGG pathways for the MM-exclusive genes included *non-small cell lung cancer*, *endometrial cancer*, *Epstein-Barr virus (EBV) infection, base excision repair* and *mismatch repair*-which are pointers to DNA damage and increased risk of cancer development - and also pathways for *Parkinson’s* and *prion disease* thus reflecting distinct neurodegenerative gene expression patterns in MM.

### Cancer, DNA repair and PD pathways are associated with MM

To identify pathways associated with the different states of malaria, GO and KEGG pathway enrichment analyses were conducted. The most significant pathways that correlated with the 352 genes specific to MM (Fig. 2f, suppl. Table 1g) were *Non-small cell lung cancer* (p< 0.0007), *Endometrial cancer* (0.0016), *Epstein-Barr virus (EBV) infection* (0.0046), *DNA replication* (p< 0.0010), *Base excision repair* (0.0055) and *Mismatch repair* (p< 0.0132). These pathways are suggestive of damage to host DNA and the subsequent increased risk of cancer development. Interestingly, the pathways for *Parkinson’s disease* (PD) and *Prion disease* (PrD) were up-regulated (Fig. 2f, suppl. Table 1g). This shows that MM though mild, could have deleterious implications on the brain.

### Drug resistance, glutathione metabolism and endocannabinoid signaling correlate with NCM and CM

The most significant pathways that were associated with NCM (suppl. Table 1f) included *Platinum drug resistance* (p< 0.0025) *Nucleotide excision repair* (p< 0.0226) and *Glutathione metabolism* (p< 0.0373), which implies drug resistance, DNA damage and antioxidant processes during *P. falciparum* infection. With regards to CM (Fig. 2e, suppl. Table 1e), pathways involving *Retrograde endocannabinoid signaling* (p< 0.0265) and *Non-alcoholic fatty liver disease (NAFLD)* (p< 0.0269) were most relevant. This could reflect an endocannabinoid modulation in CM pathology and hepatocyte injury.

### GO analysis of genes enriched in MM

We subsequently assessed the GOs elevated in *P. falciparum* infection. Fig. 2d and suppl. Table 1d display a selection of significant GOs from all three categories Biological Process (BP), Cellular Component (CC) and Molecular Function (MF) in MM. For the GO-BPs the terms *biogenesis*, *regulation of nitrogen compound metabolic process, positive regulation of protein metabolic process, response to biotic stimulus, DNA damage response, detection of DNA damage, positive regulation of phospholipid metabolic process*, *blood vessel remodeling, cell cycle arrest, response to unfolded protein, immune response* and *interleukin (IL)-10 production* were significant. These are indicative of the host repair and recovery mechanisms following macromolecular damage and inflammation. Amongst the GO-CCs, the terms *cytoplasm, intracellular, nucleoplasm* and *membrane-bounded organelle* were the most significant. In the GO-MFs, the major terms that came up were *RNA polymerase II distal enhancer sequence-specific DNA binding, oxidized purine DNA binding, mismatch repair complex binding* and *damaged DNA binding*, which are indications of transcriptional processes and DNA repair mechanisms in the host [44].

### Down-regulation of cytoskeletal organization, neutrophil degranulation, aberrant signaling associated GOs are enriched in NCM and CM

Furthermore, we evaluated the GO terms that are over-represented in NCM and CM (Fig. 2a, b, suppl. Table 1b, c). For GO-BPs, the terms *microtubule organizing center organization, glutathione metabolic process, glutathione derivative biosynthetic process, oxidation-reduction process, DNA damage checkpoint, mitotic cell cycle arrest, reactive oxygen species biosynthetic process, removal of superoxide radicals, detoxification, activation of mitogen-activated protein* kinases *(MAP3Ks) activity, negative regulation of cytoskeleton organization and amyloid-beta clearance* were significant. These terms are suggestive of *P. falciparum* mediated antioxidant synthesis and depletion, cell cycle arrest due to DNA damage, cytoskeletal destabilization and possibly, neurological impairment. With regards to the GO-CCs the terms *intracellular, cytoplasm, membrane-enclosed lumen, cell*, and *transcriptional repressor complex* were noteworthy. Significant in the GO-MFs were the terms *superoxide dismutase copper chaperone activity, glutathione transferase activity, class I DNA-(apurinic or apyrimidinic site) endonuclease activity* and *protein kinase activity*. These terms are pointers to host defense mechanisms against *falciparum* infection by up-regulating anti-oxidative activities and DNA damage repair. It also signifies a probable increase in protein kinases activity which may destabilize microtubule organization [45, 46].

The GO-BPs that were significant in CM includes *signal transduction in absence of ligand, neutrophil degranulation, negative regulation of gene expression, neutrophil activation* and *extrinsic apoptotic signaling pathway.* These terms are indicative of parasite driven abnormal signaling and inflammatory provoked tissue destruction in CM [47, 48]. In the GO-CCs, *tertiary granule membrane specific granule membrane intracellular organelle part chromosome* and *centromeric region part* were the most significant terms. The terms highly represented in GO-MF for CM were *oxidoreductase activity, acting on paired donors, with incorporation or reduction of molecular oxygen*, *proximal promoter DNA-binding transcription repressor activity, RNA polymerase II-specific* and *zinc ion binding* which demonstrate host defense against ROS, transcription suppression and the necessity of zinc in binding of the *P. falciparum* to host cells [49].

### Differential expression between CM and NCM

We set out to further refine our results from the Venn diagram analysis of gene expression by investigating the differences between CM and NCM. Fig. 3 shows signatures of differentially up- and down-regulated genes and associated GOs in CM compared to NCM (suppl. Table 2a). For the genes up-regulated between CM and NCM NF-kappaB signaling, immune response, response to herbicide and neutrophil activation were amongst the most significantly elevated GO terms (Fig. 3a, suppl. Table 2b). For the genes down-regulated between CM and NCM myeloid cell homeostasis, regulation of axon diameter, hemopoiesis were amongst the most significantly elevated GO terms (Fig. 3b, suppl. Table 2c). The heatmap and hierarchical cluster analysis in Fig. 3c displays a cluster with higher expression in CM (red) and a cluster with lower expression in CM (green). In Fig. 3d, the signatures of the genes up-regulated (ratio > 2, p < 0.05) and down-regulated (ratio < 0.5, p < 0.05) between CM and NCM are listed. Interestingly, IL-18 receptor 1 (*IL18R1*) is amongst the up-regulated genes indicating the involvement of IL-18 signaling.

**Figure 3.**
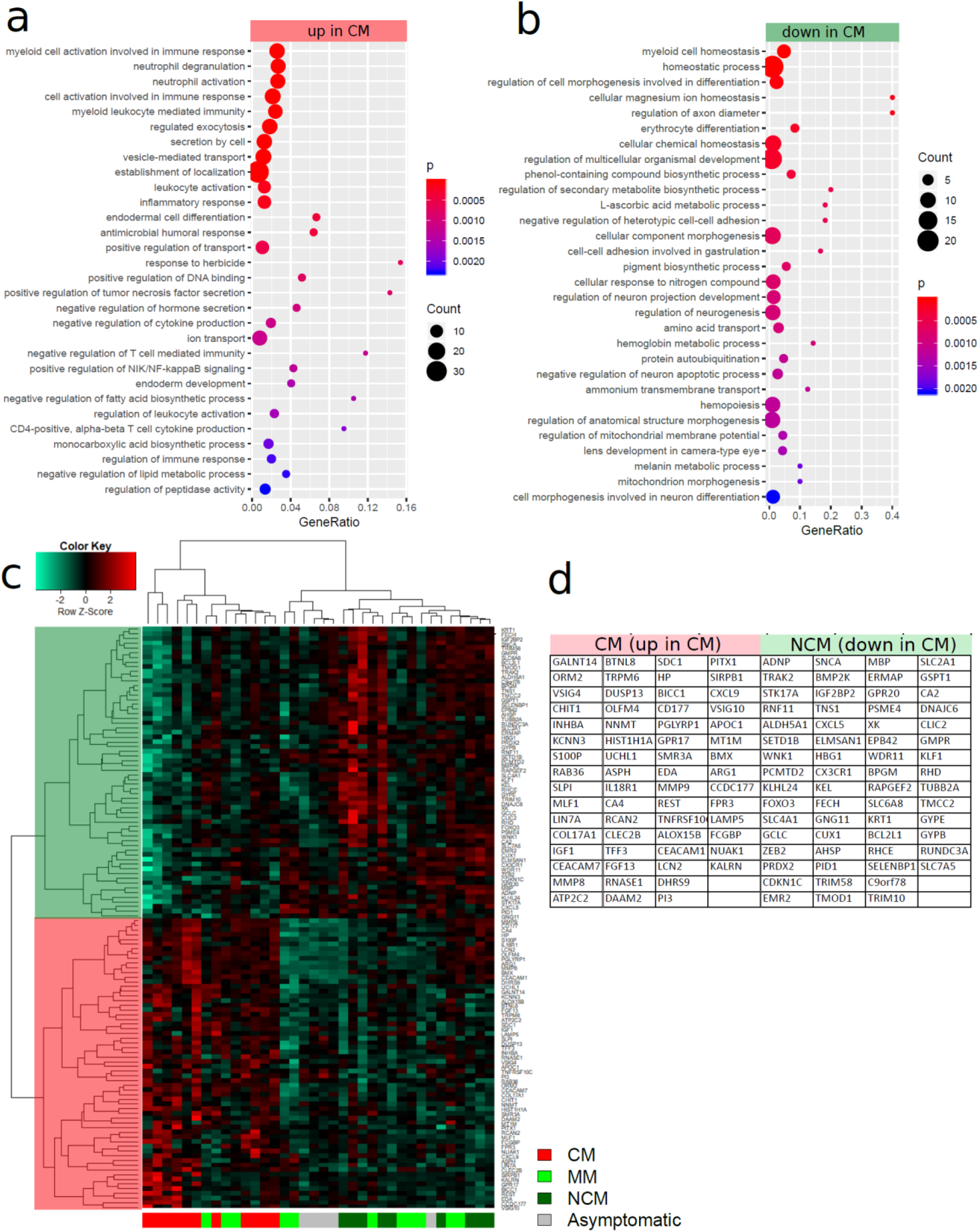
Differential expression analysis between CM and NCM reveals NF-kappaB signaling, immune response, response to herbicide and neutrophil activation amongst GO terms overrepresented in up-regulated genes whereas myeloid cell homeostasis, regulation of axon diameter, hemopoiesis were amongst GO terms highly expressed in downregulated genes. (a) Most significant GO terms elevated in genes up-regulated between CM and NCM. (b) Most significant GO terms elevated in genes down-regulated between CM and NCM. (c) Hierarchical cluster analysis and heatmap of genes up-regulated (ratio > 2, p < 0.05) and down-regulated (ratio < 0.5, p < 0.05) between CM and NCM. The cluster of genes up-regulated in CM is marked with red shading, the cluster of down-regulated genes with green shading. The genes associated with both clusters are listed in (d). IL18R1 among the up-regulated genes is indicative of the involvement of IL18-signaling.

### Protein interaction network and regulating transcription factors

For further elucidation of protein interaction networks and transcription factors involved in gene regulatory networks constituting CM we used Metascape analysis of genes differentially expressed between CM and NCM (Fig. 4). Fig. 4a shows the protein interaction network which besides the IL-18 parts already mentioned, has a large connected part centered around hexokinase 1 (HK1) and tubulin beta-2A chain (TUBB2A). Fig. 4b displays the results of the Metascape TRRUST analysis[43] indicating involvement of the transcription factors IRF1, STAT1/3, NFKB1 and RELA in the gene regulatory networks of the CM state.

**Figure 4.**
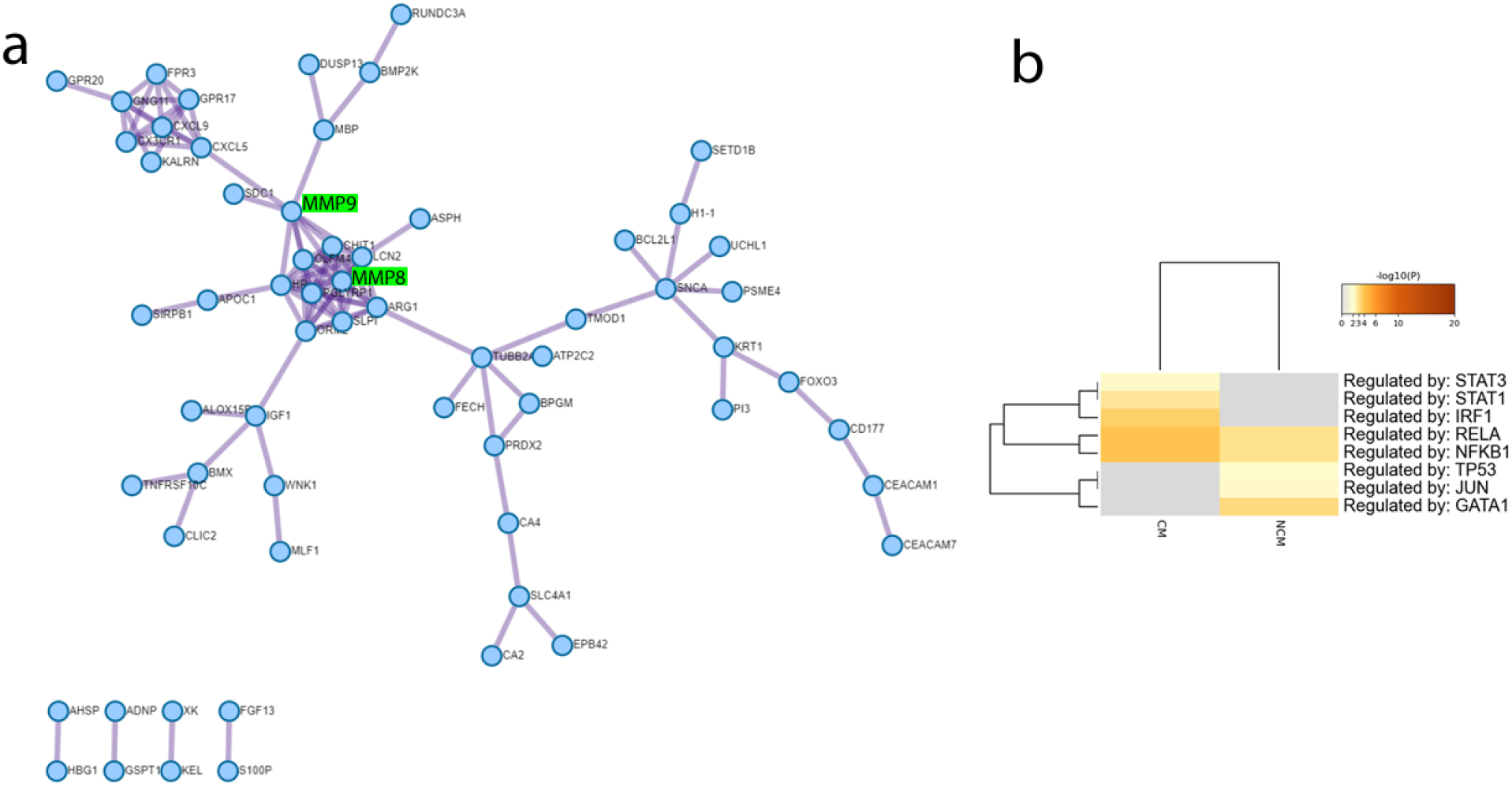
Metascape analysis of genes differentially expressed between CM and NCM reveals protein interaction network and involvement of the transcription factors IRF1, STAT1/3, NFKB1 and RELA in the gene regulatory networks of the CM state. (a) Protein interaction network generated with Metascape from genes differentially expressed between CM and NCM. A central cluster is located around MMP8 and MMP8 which have been associated with CM by Polimeni and Prato, due to their potential to disrupt the subendothelial basement membrane and process and regulate cytokines [16]. Moreover, Polimeni and Prato propose that hemozoin promotes NF-kappaB-controlled gene expression of MMP9. (b) Heatmap of transcription factors generated with Metascape employing the TRRUST method using genes differentially expressed between CM and NCM. Graphics were created with Metascape (https://metascape.org/), [42].

## Discussion

In this analysis, pooled transcriptome data from whole blood cells of malaria patients which involved MM, NCM and CM were evaluated. Although transcriptome organization is reported to be poorly preserved between brain and blood, there are some brain co-expression modules which demonstrate strong evidence of preservation in blood and could be useful for identifying blood biomarkers for neurological diseases [50]. Therefore, some extrapolations can be made from transcriptome data from blood (which is more readily accessible) to deduce genes and pathways associated with neurological conditions. Previous investigations by Thiam et al. and Boldt et al., which we employed in this analysis had successfully used blood transcriptional analysis to detect changes in gene expression in CM, NCM and MM. In this study, it was discovered from the cluster analysis that the 3 states of *P. falciparum* infection have some common characteristics between them although certain CM states could be distinctly different. While CM was mainly associated with NCM, there was some clustering between MM and CM. This might be an indication that malaria caused by *P. falciparum*, whether mild or severe begins with a more generic effect on the host. The clinical manifestation of which is described as fever, malaise, headache, nausea and vomiting, though MM could rapidly progress into CM if not treated quickly [51, 52]. Furthermore, a study by Taylor et al. showed that the classification of CM in children is sometimes misdiagnosed, where coma has other causes and parasitemia is incidental [53]. This suggests that the detection of CM-specific characteristics can be challenging, which could result in its clustering with the other states of *falciparum* malaria.

In addition, we went on to identify genes that are expressed in the 3 *falciparum* induced conditions. It was discovered that 2876 genes were expressed in common between them and CM surprisingly had the least number of expressed genes. In all, genes associated with host response to parasite invasion, inflammatory response, cellular stress, detoxification of toxic metals and clathrin mediated vesicle assembly were up-regulated. It has been established that entry of malaria parasite into the host elicit immune response which can lead to oxidative stress [54]. Oxidative stress caused by the release of parasite toxins such as HMZ or inflammatory response to *falciparum* entry have been implicated in cellular stress, especially in severe malaria [5, 55]. Moreover, levels of metals such as copper and zinc are increased in acute malaria, which could be as a result of the host’s inflammatory response against parasite and lysis of erythrocytes [56]. Concurrently, there is trafficking of the virulent protein, *P. falciparum* erythrocyte membrane protein 1 (PfEMP1) to the surface of infected red blood cells (iRBCs), which involves membranous organelles termed Maurer’s clefts [57]. Clathrin and its adaptor proteins possibly participate in the process hence its upregulation in the 3 disease states [58]. The genes connected to mental retardation and AD were also present in all 3 of *falciparum* malaria. Previous investigations have documented the predisposition of children with CM to develop mental disorders probably as a result of ischemic and hypoxic neural injury[59, 60]. The elevation of the expression levels of these genes in MM and NCM implies the possible detrimental nature of these *falciparum* infections on the brain as well. Indeed, some studies have documented cognitive deficits in children and adults who suffered from acute MM and NCM [61–63]. Using an animal model, Guha et al. proved that a single incident of MM induced microglial activation, neuroinflammation and behavioral changes accompanied by increase in pro-inflammatory cytokine expression in the brain [64]. Hence, the severity of neurological deficits reported in CM could be attributed to the down-regulation of various genes involved in cell survival, protein synthesis and mitochondrial respiration. Additionally, a study by Bodlt et al. discovered the down-regulation of a number of genes in CM of Gabonese children due to expression of hypoxia induced genes aryl hydrocarbon receptor (AhRF), GA binding protein transcription factor (GABP) and hypoxia inducible factor-1 (HIF1) [30]. Brain hypoxia is observed in CM owing to sequestration of infected erythrocytes, inflammatory cytokine production, vessel leakage and occlusion [65].

With regards to pathways associated with the different *falciparum* malaria, KEGG enrichment analysis unveiled the association of MM with the neurodegenerative diseases, PD and PrD. This means MM and some neurodegenerative diseases may share parallel biological processes, which is contradictory to the findings of Thiam et al. and Cabantous et al., where neurodegenerative pathways were found in CM only. Nonetheless, processes involved in cancer, EBV infection and DNA damage repair were the most significant. Some human pathogens including *falciparum* and EBV have been implicated as etiologic agents in the development of cancer [66]. The invasion of *falciparum* and subsequent release of HMZ into the host induces inflammation which facilitates ROS production and cellular stress [67, 68]. ROS production even at low levels can lead to gene and chromosomal mutations via DNA double strand breaks (DSBs), which is one of the most severe forms of DNA damage [69]. DSBs when unrepaired or mis-repaired, can cause stress, cell death, chromosome instability, and cancer [70]. EBV infection is also associated with ROS generation [71, 72] and since cancer development can stem from DNA damage, it is not surprising that these pathways were elevated in MM. Again, DNA damage and DNA damage response are hallmarks of neurodegenerative diseases including PD and PrD [73]. The up-regulation of the neurodegenerative pathways in MM suggests a shared cytotoxic mechanism between these diseases and buttresses earlier observation that MM could have implications on brain function.

Similarly, KEGG pathway enrichment assessment was done for NCM and CM. Platinum drug resistance, nucleotide excision repair and glutathione metabolism were associated with NCM. Resistance to antimalarial drugs is a growing challenge in malarial treatment [74]. The up-regulation of a drug resistance pathway similar to that observed in cancer during NCM implies a parasite facilitated host resistance to antimalarials. This might be as a result of host cellular modification especially with DNA damage and repair mechanisms which were also up-regulated in NCM. For instance, multiple efflux and uptake transporters are present in red blood cells and hepatocytes [75, 76]. Increase in and genetically altered drug efflux transporters reduces antimalarial drug uptake and concentrations in red blood cells [77]. This ensues in minimum effective drug concentration, inhibiting the termination of malaria infection and potentially, the development of resistance. The association of glutathione metabolism in NCM is an indication of glutathione modulation of disease progression particularly with regards to malarial treatment. A study by Zuluaga et al. revealed that amodiaquine treatment failure was connected to erythrocytic glutathione [78]. This is because glutathione competes with amodiaquine for the heme group, which could result in therapeutic failure [78]. Again, the concentration of glutathione was decreased in iRBCs, which is an indication of the antioxidant consumption due to ROS generation [79]. Thus, glutathione though an important antioxidant defense in malaria may likely modulate the host’s response to drug treatment. The pathways that correlated with CM were retrograde endocannabinoid signaling and NAFLD. Previous researches have described the up-regulation of certain genes involved in neurodegenerative diseases including amyloid precursor protein (*APP)*, ubiquilins *(UBQLN)* and Jun proto-oncogene *(JUN)* as candidate genes for NAFLD [80–83]. In their study, Karbalaei et al. showed that there were 190 genes expressed in common between NAFLD and AD and that NAFLD has an undoubted relation to AD [83]. Moreover, the endocannabinoids and their receptors (especially cannabinoid 1, CB1) are known to induce steatosis and lipogenic gene expression resulting in NAFLD [84, 85]. The cannabinoid 2 (CB2) receptor on the other hand regulates neuro-inflammatory responses and affects various macrophage functions, including antigen uptake and presentation and chemokine/cytokine production [86–89]. A study by Alferink et al. reported that mice with a deletion of the CB2-encoding gene (Cnr2^−^/^−^) and immunized with *Plasmodium berghei* ANKA erythrocytes showed enhanced survival and diminished BBB disruption in experimental CM [86]. Furthermore, the CB1 receptor antagonist rimonabant, is reported to have neuroprotective properties in some animal models of neurodegenerative disorders [90]. In addition, endocannabinoid molecules were significantly increased in the acute phase of *falciparum* infection in children [91]. Together, these results reveal that genes involved in NAFLD and the endocannabinoid system could serve as potential biomarkers for malaria severity. This could also be an indication that the consequence of CM on the brain may comprise some neurological pathways other than those classically described for neurodegenerative diseases.

Additionally, we assessed the GOs up-regulated in the 3 *falciparum* infections and in MM analysis showed biological processes related to biogenesis, host metabolic processes, DNA damage repair activities and anti-inflammation with the up-regulation of IL-10. These terms corroborate reports that *falciparum* invasion causes damage to host molecules and inflammation. In MM however, there seem to be an increase in metabolic and damage control mechanisms to counteract the negative effects of ROS and inflammatory products.

The GOs elevated in NCM revealed destabilization of cellular cytoskeleton, ROS production, DNA damage checkpoint and amyloid-beta clearance. The surge in ROS levels during oxidative stress is implicated in the increase in the activation of MAP3Ks pathways [92]. An aberration from the strict control of MAP3Ks signaling is connected to the development of many neurodegenerative diseases including AD and PD [93]. Activated MAP3K signaling pathways are thought to contribute to neurodegenerative pathogenesis through the phosphorylation of APP and α-synuclein, leading to aggregates formation that trigger neuronal apoptosis [93–95]. Again, an increase in MAPKs signaling induces cytoskeletal abnormalities through unusual phosphorylation and consequent aggregation of cytoskeletal elements [96–98]. For example, the hyper-phosphorylation of the microtubule associated protein, tau results in its aggregation, microtubule destabilization and the formation of neurofibrillary tangles with subsequent degeneration of neurons [99–102]. Also, axonal injury with the accumulation of APP has been reported in post-mortem studies of CM patients, both adults and pediatric cases [103–105]. Accordingly, the amyloid-beta clearance observed in NCM could represent a reversible pathway to neurological damage in NCM. With respect to CM, abnormal signaling, neutrophil degranulation, repression of gene expression and apoptosis were the predominant GO terms. Using an experimental model of CM, a recent study by Kumar et al. reported an increase in D1 and D2 dopaminergic receptor expression, phosphorylated dopamine- and cAMP-regulated phosphoprotein (DARPP), p25, cyclin-dependent kinase 5 (CDK5), Ca^2+^/calmodulin-dependent protein kinase IIα (CaMKIIα) and D1-D2 heteromers [106]. The dysregulation of the dopaminergic receptors led to the impairment and degeneration of medium spiny neurons [106]. Findings from neuroimaging studies have demonstrated that the anatomical and functional grouping of dopaminergic neurons in striatum performs a significant role in the execution of cognition and behavioral consequences in young adults [107, 108]. The cognitive deficit observed in CM survivors therefore could be attributed in part to the dysregulation of the dopaminergic system. Although neutrophils are scarce in the central nervous under normal physiological conditions, they infiltrate the brain in several pathological conditions including malaria [109]. The accumulation of these granulocytes lead to the release of neutrophil extracellular traps (NETs) which directly damage the BBB, surrounding neurons and consequently apoptosis [48]. Boldt et al. in their findings showed a down-regulation of genes in CM patients [30]. Hence, the up-regulation of genes involved in gene repression is an indicator that several genes involved in cellular metabolism and function are down-regulated in CM, which accounts for the severity of the disease as compared to MM.

Most outstanding of the inflammatory processes found associated with the genes up-regulated in CM were neutrophil activation and IL18-signalling. Previous publications have described IL18 as a major player in the pathogenesis of severe malaria through a pathway of elevating IFN-gamma [110]. The activation of this pathway is further supported by the transcription factor Interferon regulatory factor 1 (IRF1) for which we found a regulatory role in CM via Metascape analysis. The relevance of neutrophils for inflammatory aggravation of malaria has been described by Knacksted et al. [111]. This is mediated by the extrusion of chromatin in the form of NETs upon neutrophil death. In addition, up-regulation of genes linked to herbicide response reiterate the significance of the “plant-like” apicoplast in *falciparum* virulence [112]. The GO term *regulation of axon diameter* found in the down-regulated genes in CM provides a mechanistic clue of neurological impairments in CM. The involved genes *XK* and *KEL* were furthermore reported to induce neuropathological abnormalities such as giant axons when knocked out in mice [113]. The protein interaction network besides relevant sub-networks of IL-18 signaling and the blood group proteins XK and KEL regulating axon diameter showed a large connected area centered by TUBB2A and the Metalloproteinases 8 and 9 (MMP8, MMP9). TUBB2A has been reported to be up-regulated in CM by Cabantous et al. [29] as part of the Parkin-Ubiquitin proteasome degradation pathway which when impaired can be deleterious to dopaminergic neurons [114]. MMP8 and MMP9 which is activated by HMZ have been associated with CM by Polimeni and Prato et al. due to their potential to disrupt the subendothelial basement membrane and process and regulate cytokines [16].

A limitation of this study is that, the number of patients enrolled in the investigations were few and hence larger studies are required to determine the pernicious effect of *falciparum* infection on the brain. Furthermore, the study was based only on blood-derived cells and included no brain biopsies thus implying that some neurological sequelae are already manifested in the genetic networks in blood-derived cells

## Conclusion

We have shown in this study that all forms of malaria including MM could have detrimental implications on the brain. The difference in the severity of the various forms of malaria is however dependent on which genes are down/up-regulated in its progression. Also, given the up-regulation of the endocannabinoid and the dysregulation of dopaminergic systems reported in CM, studies into the interplay of CB1/CB2 and D1/D2 receptors and their involvement in neurological sequelae may be conducted. Howbeit, the apicoplast remains a vital target for antimalarial drugs. We anticipate that our data might form the basis for future hypothesis-driven malaria research.

## List of abbreviations

AhRF: Aryl hydrocarbon receptor
AD: Alzheimer’s disease
APP: Amyloid precursor protein
BBB: Blood brain barrier
BP: Biological Process
CaMKIIα: Ca^2+^/calmodulin-dependent protein kinase IIα
CB1/CB2: Cannabinoid receptors 1/2
CC: Cellular Component
CM: Cerebral malaria
CDK5: Cyclin-dependent kinase 5
D1/D2: Copaminergic receptors 1/2
DARPP: Dopamine- and cAMP-regulated phosphoprotein
DSBs: Double strand breaks
EBV: Epstein-Barr virus
GABP: GA binding protein transcription factor
GO: Gene ontology
HK1: Hexokinase 1
HMZ: Hemozoin
HIF1: Hypoxia inducible factor-1
iRBCs: Infected red blood cells
IRF1: Interferon Regulatory Factor 1
JUN: Jun proto-oncogene
KEL: Kell metallo-endopeptidase
KEGG: Kyoto Encyclopedia of Genes and Genomes
MAP3Ks: mitogen-activated protein kinases
MF: Molecular Function
MM: mild malaria
MMP8/9: Metalloproteinases 8 and 9
NAFLD: non-alcoholic fatty liver disease
NCM: non-cerebral severe malaria
NET: Neutrophil extracellular traps
NFKB1: Nuclear Factor Kappa B Subunit 1
PfEMP1: *Plasmodium falciparum* erythrocyte membrane protein 1
PD: Parkinson’s disease
PrD: Prion disease
RELA: RELA proto-oncogene, NF-KB Subunit
ROS: Reactive oxygen species
STAT1/3: Signal transducer and activator of transcription 1/3
TUBB2A: Tubulin beta-2A chain
UBQLN: Ubiquilins
WHO: World health organization
XK: Membrane transport protein X-linked-Kx

## Declarations

### Ethics approval and consent to participate

Not applicable.

### Consent for publication

Not applicable.

### Availability of data and materials

The datasets analysed during the current study are available in the National Center for Biotechnology Information - Gene Expression Omnibus (NCBI GEO) repository under the accessions GSE116306 (https://www.ncbi.nlm.nih.gov/geo/query/acc.cgi?acc=GSE116306) and GSE1124 (https://www.ncbi.nlm.nih.gov/geo/query/acc.cgi?acc=GSE1124). The datasets are summarized in Table 1.

### Authors’ contributions

A.A.K., W.W. and J.A. wrote the manuscript. W.W. performed the pooled analysis and follow-up analyses. J.A. and A.A.K. initiated and conceived this study. A.A.K. and W.W. contributed equally to this work.

## Acknowledgements

J.A. acknowledges the medical faculty of Heinrich Heine University for financial support.

## References

1. https://www.who.int/publications/i/item/9789240015791: World malaria report 2020: 20 years of global progress and challenges. 30.11.2020 edition. Geneva: World Health Organization; 2020.

2. Mukhtar MM, Eisawi OA, Amanfo SA, Elamin EM, Imam ZS, Osman FM, Hamed ME: Plasmodium vivax cerebral malaria in an adult patient in Sudan. Malaria Journal 2019, 18: 316.

3. Eugenin EA, Martiney JA, Berman JW: The malaria toxin hemozoin induces apoptosis in human neurons and astrocytes: Potential role in the pathogenesis of cerebral malaria. Brain research 2019, 1720: 146317–146317.

4. Imai T, Iwawaki T, Akai R, Suzue K, Hirai M, Taniguchi T, Okada H, Hisaeda H: Evaluating experimental cerebral malaria using oxidative stress indicator OKD48 mice. Int J Parasitol 2014, 44: 681–685.

5. Kavishe RA, Koenderink JB, Alifrangis M: Oxidative stress in malaria and artemisinin combination therapy: Pros and Cons. The FEBS Journal 2017, 284: 2579–2591.

6. Simão-Gurge RM, Wunderlich G, Cricco JA, Cubillos EFG, Doménech-Carbó A, Cebrián-Torrejón G, Almeida FG, Cirulli BA, Katzin AM: Biosynthesis of heme O in intraerythrocytic stages of Plasmodium falciparum and potential inhibitors of this pathway. Scientific Reports 2019, 9: 19261.

7. Nagaraj VA, Sundaram B, Varadarajan NM, Subramani PA, Kalappa DM, Ghosh SK, Padmanaban G: Malaria Parasite-Synthesized Heme Is Essential in the Mosquito and Liver Stages and Complements Host Heme in the Blood Stages of Infection. PLOS Pathogens 2013, 9: e1003522.

8. Lim L, McFadden GI: The evolution, metabolism and functions of the apicoplast. Philosophical transactions of the Royal Society of London Series B, Biological sciences 2010, 365: 749–763.

9. Pishchany G, Skaar EP: Taste for blood: hemoglobin as a nutrient source for pathogens. PLoS pathogens 2012, 8: e1002535–e1002535.

10. Slater AF, Cerami A: Inhibition by chloroquine of a novel haem polymerase enzyme activity in malaria trophozoites. Nature 1992, 355: 167–169.

11. Grau GE, Mackenzie CD, Carr RA, Redard M, Pizzolato G, Allasia C, Cataldo C, Taylor TE, Molyneux ME: Platelet accumulation in brain microvessels in fatal pediatric cerebral malaria. J Infect Dis 2003, 187: 461–466.

12. Sullivan AD, Ittarat I, Meshnick SR: Patterns of haemozoin accumulation in tissue. Parasitology 1996, 112 (Pt 3): 285–294.

13. Kwiatkowski D, Bate CA, Scragg IG, Beattie P, Udalova I, Knight JC: The malarial fever response--pathogenesis, polymorphism and prospects for intervention. Ann Trop Med Parasitol 1997, 91: 533–542.

14. Sherry BA, Alava G, Tracey KJ, Martiney J, Cerami A, Slater AF: Malaria-specific metabolite hemozoin mediates the release of several potent endogenous pyrogens (TNF, MIP-1 alpha, and MIP-1 beta) in vitro, and altered thermoregulation in vivo. J Inflamm 1995, 45: 85–96.

15. Ihekwereme CP, Esimone CO, Nwanegbo EC: Hemozoin inhibition and control of clinical malaria. Advances in pharmacological sciences 2014, 2014: 984150–984150.

16. Polimeni M, Prato M: Host matrix metalloproteinases in cerebral malaria: new kids on the block against blood-brain barrier integrity? Fluids and barriers of the CNS 2014, 11: 1–1.

17. Oluwayemi IO, Brown BJ, Oyedeji OA, Oluwayemi MA: Neurological sequelae in survivors of cerebral malaria. Pan Afr Med J 2013, 15: 88.

18. Bhandary B, Marahatta A, Kim H-R, Chae H-J: An Involvement of Oxidative Stress in Endoplasmic Reticulum Stress and Its Associated Diseases. International Journal of Molecular Sciences 2013, 14: 434–456.

19. Hussain T, Tan B, Yin Y, Blachier F, Tossou MCB, Rahu N: Oxidative Stress and Inflammation: What Polyphenols Can Do for Us? Oxidative Medicine and Cellular Longevity 2016, 2016: 7432797.

20. Muthuswamy AD, Vedagiri K, Ganesan M, Chinnakannu P: Oxidative stress-mediated macromolecular damage and dwindle in antioxidant status in aged rat brain regions: role of L-carnitine and DL-alpha-lipoic acid. Clin Chim Acta 2006, 368: 84–92.

21. Kannan K, Jain SK: Oxidative stress and apoptosis. Pathophysiology 2000, 7: 153–163.

22. Redza-Dutordoir M, Averill-Bates DA: Activation of apoptosis signalling pathways by reactive oxygen species. Biochimica et Biophysica Acta (BBA) - Molecular Cell Research 2016, 1863: 2977–2992.

23. Hegde ML, Hegde PM, Rao KS, Mitra S: Oxidative genome damage and its repair in neurodegenerative diseases: function of transition metals as a double-edged sword. Journal of Alzheimer’s disease : JAD 2011, 24 Suppl 2: 183–198.

24. Poetsch AR: The genomics of oxidative DNA damage, repair, and resulting mutagenesis. Computational and Structural Biotechnology Journal 2020, 18: 207–219.

25. Sertan Copoglu U, Virit O, Hanifi Kokacya M, Orkmez M, Bulbul F, Binnur Erbagci A, Semiz M, Alpak G, Unal A, Ari M, Savas HA: Increased oxidative stress and oxidative DNA damage in non-remission schizophrenia patients. Psychiatry research 2015, 229: 200–205.

26. Singh A, Kukreti R, Saso L, Kukreti S: Oxidative Stress: A Key Modulator in Neurodegenerative Diseases. Molecules 2019, 24.

27. Li J, Shang Y, Wang L, Zhao B, Sun C, Li J, Liu S, Li C, Tang M, Meng F-L, Zheng P: Genome integrity and neurogenesis of postnatal hippocampal neural stem/progenitor cells require a unique regulator Filia. Science Advances 2020, 6: eaba0682.

28. Thiam A, Sanka M, Ndiaye Diallo R, Torres M, Mbengue B, Nunez NF, Thiam F, Diop G, Victorero G, Nguyen C, et al: Gene expression profiling in blood from cerebral malaria patients and mild malaria patients living in Senegal. BMC Medical Genomics 2019, 12: 148.

29. Cabantous S, Poudiougou B, Bergon A, Barry A, Oumar AA, Traore AM, Chevillard C, Doumbo O, Dessein A, Marquet S: Understanding Human Cerebral Malaria through a Blood Transcriptomic Signature: Evidences for Erythrocyte Alteration, Immune/Inflammatory Dysregulation, and Brain Dysfunction. Mediators of Inflammation 2020, 2020: 3280689.

30. Boldt ABW, van Tong H, Grobusch MP, Kalmbach Y, Dzeing Ella A, Kombila M, Meyer CG, Kun JFJ, Kremsner PG, Velavan TP: The blood transcriptome of childhood malaria. EBioMedicine 2019, 40: 614–625.

31. Gentleman RC, Carey VJ, Bates DM, Bolstad B, Dettling M, Dudoit S, Ellis B, Gautier L, Ge Y, Gentry J, et al: Bioconductor: open software development for computational biology and bioinformatics. Genome Biol 2004, 5: R80.

32. Smyth GK: Linear models and empirical bayes methods for assessing differential expression in microarray experiments. Stat Appl Genet Mol Biol 2004, 3: Article3.

33. Gautier L, Cope L, Bolstad BM, Irizarry RA: affy--analysis of Affymetrix GeneChip data at the probe level. Bioinformatics 2004, 20: 307–315.

34. Johnson WE, Li C, Rabinovic A: Adjusting batch effects in microarray expression data using empirical Bayes methods. Biostatistics 2007, 8: 118–127.

35. Leek JT, Johnson WE, Parker HS, Jaffe AE, Storey JD: The sva package for removing batch effects and other unwanted variation in high-throughput experiments. Bioinformatics 2012, 28: 882–883.

36. Storey JD: A direct approach to false discovery rates. Journal of the Royal Statistical Society: Series B (Statistical Methodology) 2002, 64: 479–498.

37. Warnes GR, Bolker B, Bonebakker L, Gentleman R, Liaw WHA, Lumley T, Maechler M, Magnusson A, Moeller S, Schwartz M: Gplots: various R programming tools for plotting data. 2016. R package version 2014, 2.

38. Chen H, Boutros PC: VennDiagram: a package for the generation of highly-customizable Venn and Euler diagrams in R. BMC Bioinformatics 2011, 12: 35.

39. Kanehisa M, Furumichi M, Tanabe M, Sato Y, Morishima K: KEGG: new perspectives on genomes, pathways, diseases and drugs. Nucleic Acids Res 2017, 45: D353–d361.

40. Falcon S, Gentleman R: Using GOstats to test gene lists for GO term association. Bioinformatics 2007, 23: 257–258.

41. Wickham H: ggplot2: Elegant Graphics for Data Analysis. Springer Publishing Company, Incorporated; 2009.

42. Zhou Y, Zhou B, Pache L, Chang M, Khodabakhshi AH, Tanaseichuk O, Benner C, Chanda SK: Metascape provides a biologist-oriented resource for the analysis of systems-level datasets. Nature Communications 2019, 10: 1523.

43. Han H, Cho JW, Lee S, Yun A, Kim H, Bae D, Yang S, Kim CY, Lee M, Kim E, et al: TRRUST v2: an expanded reference database of human and mouse transcriptional regulatory interactions. Nucleic Acids Res 2018, 46: D380–d386.

44. Sampaio NG, Emery SJ, Garnham AL, Tan QY, Sisquella X, Pimentel MA, Jex AR, Regev-Rudzki N, Schofield L, Eriksson EM: Extracellular vesicles from early stage Plasmodium falciparum-infected red blood cells contain PfEMP1 and induce transcriptional changes in human monocytes. Cell Microbiol 2018, 20: e12822.

45. Cavallini A, Brewerton S, Bell A, Sargent S, Glover S, Hardy C, Moore R, Calley J, Ramachandran D, Poidinger M, et al: An Unbiased Approach to Identifying Tau Kinases That Phosphorylate Tau at Sites Associated with Alzheimer Disease. Journal of Biological Chemistry 2013, 288: 23331–23347.

46. Morshed N, Lee M, Rodriguez FH, Lauffenburger DA, Mastroeni D, White F: Quantitative phosphoproteomics uncovers dysregulated kinase networks in Alzheimer’s disease. bioRxiv 2020: 2020.2008.2018.255778.

47. Knackstedt SL, Georgiadou A, Apel F, Abu-Abed U, Moxon CA, Cunnington AJ, Raupach B, Cunningham D, Langhorne J, Krüger R, et al: Neutrophil extracellular traps drive inflammatory pathogenesis in malaria. Science immunology 2019, 4: eaaw0336.

48. Boeltz S, Muñoz LE, Fuchs TA, Herrmann M: Neutrophil Extracellular Traps Open the Pandora’s Box in Severe Malaria. Front Immunol 2017, 8: 874.

49. Marvin RG, Wolford JL, Kidd MJ, Murphy S, Ward J, Que EL, Mayer ML, Penner-Hahn JE, Haldar K, O’Halloran TV: Fluxes in “free” and total zinc are essential for progression of intraerythrocytic stages of Plasmodium falciparum. Chemistry & biology 2012, 19: 731–741.

50. Cai C, Langfelder P, Fuller TF, Oldham MC, Luo R, van den Berg LH, Ophoff RA, Horvath S: Is human blood a good surrogate for brain tissue in transcriptional studies? BMC Genomics 2010, 11: 589.

51. Bartoloni A, Zammarchi L: Clinical aspects of uncomplicated and severe malaria. Mediterranean journal of hematology and infectious diseases 2012, 4: e2012026–e2012026.

52. Marrelli MT, Brotto M: The effect of malaria and anti-malarial drugs on skeletal and cardiac muscles. Malaria Journal 2016, 15: 524.

53. Taylor TE, Fu WJ, Carr RA, Whitten RO, Mueller JS, Fosiko NG, Lewallen S, Liomba NG, Molyneux ME: Differentiating the pathologies of cerebral malaria by postmortem parasite counts. Nat Med 2004, 10: 143–145.

54. Long CA, Zavala F: Immune Responses in Malaria. Cold Spring Harbor perspectives in medicine 2017, 7: a025577.

55. Galluzzi L, Diotallevi A, Magnani M: Endoplasmic reticulum stress and unfolded protein response in infection by intracellular parasites. Future science OA 2017, 3: FSO198–FSO198.

56. Narsaria N, Mohanty C, Das BK, Mishra SP, Prasad R: Oxidative Stress in Children with Severe Malaria. Journal of Tropical Pediatrics 2011, 58: 147–150.

57. McHugh E, Carmo OMS, Blanch A, Looker O, Liu B, Tiash S, Andrew D, Batinovic S, Low AJY, Cho H-J, et al: Role of <span class=“named-content genus-species” id=“named-content-1”>Plasmodium falciparum</span> Protein GEXP07 in Maurer’s Cleft Morphology, Knob Architecture, and <span class=“named-content genus-species” id=“named-content-2”>P. falciparum</span> EMP1 Trafficking. mBio 2020, 11: e03320–03319.

58. Taraschi TF, Trelka D, Martinez S, Schneider T, O’Donnell ME: Vesicle-mediated trafficking of parasite proteins to the host cell cytosol and erythrocyte surface membrane in Plasmodium falciparum infected erythrocytes. International Journal for Parasitology 2001, 31: 1381–1391.

59. Idro R, Kakooza-Mwesige A, Asea B, Ssebyala K, Bangirana P, Opoka RO, Lubowa SK, Semrud-Clikeman M, John CC, Nalugya J: Cerebral malaria is associated with long-term mental health disorders: a cross sectional survey of a long-term cohort. Malaria Journal 2016, 15: 184.

60. Jain K, Sood S, Gowthamarajan K: Modulation of cerebral malaria by curcumin as an adjunctive therapy. Brazilian Journal of Infectious Diseases 2013, 17: 579–591.

61. Fernando SD, Rodrigo C, Rajapakse S: The ‘hidden’ burden of malaria: cognitive impairment following infection. Malar J 2010, 9: 366.

62. Holding PA, Snow RW: Impact of Plasmodium falciparum malaria on performance and learning: review of the evidence. Am J Trop Med Hyg 2001, 64: 68–75.

63. Kihara M, Carter JA, Newton CR: The effect of Plasmodium falciparum on cognition: a systematic review. Trop Med Int Health 2006, 11: 386–397.

64. Guha SK, Tillu R, Sood A, Patgaonkar M, Nanavaty IN, Sengupta A, Sharma S, Vaidya VA, Pathak S: Single episode of mild murine malaria induces neuroinflammation, alters microglial profile, impairs adult neurogenesis, and causes deficits in social and anxiety-like behavior. Brain Behav Immun 2014, 42: 123–137.

65. Yusuf FH, Hafiz MY, Shoaib M, Ahmed SA: Cerebral malaria: insight into pathogenesis, complications and molecular biomarkers. Infection and drug resistance 2017, 10: 57–59.

66. Robbiani Davide F, Deroubaix S, Feldhahn N, Oliveira Thiago Y, Callen E, Wang Q, Jankovic M, Silva Israel T, Rommel Philipp C, Bosque D, et al: Plasmodium Infection Promotes Genomic Instability and AID-Dependent B Cell Lymphoma. Cell 2015, 162: 727–737.

67. Percário S, Moreira DR, Gomes BAQ, Ferreira MES, Gonçalves ACM, Laurindo PSOC, Vilhena TC, Dolabela MF, Green MD: Oxidative stress in malaria. In International journal of molecular sciences, vol. 13. pp. 16346-163722012:16346–16372.

68. Keller CC, Kremsner PG, Hittner JB, Misukonis MA, Weinberg JB, Perkins DJ: Elevated Nitric Oxide Production in Children with Malarial Anemia: Hemozoin-Induced Nitric Oxide Synthase Type 2 Transcripts and Nitric Oxide in Blood Mononuclear Cells. Infection and Immunity 2004, 72: 4868–4873.

69. Sharma V, Collins LB, Chen T-H, Herr N, Takeda S, Sun W, Swenberg JA, Nakamura J: Oxidative stress at low levels can induce clustered DNA lesions leading to NHEJ mediated mutations. Oncotarget 2016, 7: 25377–25390.

70. Trenner A, Sartori AA: Harnessing DNA Double-Strand Break Repair for Cancer Treatment. Frontiers in oncology 2019, 9: 1388–1388.

71. Huang S-Y, Fang C-Y, Wu C-C, Tsai C-H, Lin S-F, Chen J-Y: Reactive Oxygen Species Mediate Epstein-Barr Virus Reactivation by N-Methyl-N’-Nitro-N-Nitrosoguanidine. PLOS ONE 2013, 8: e84919.

72. Lassoued S, Ben Ameur R, Ayadi W, Gargouri B, Ben Mansour R, Attia H: Epstein-Barr virus induces an oxidative stress during the early stages of infection in B lymphocytes, epithelial, and lymphoblastoid cell lines. Mol Cell Biochem 2008, 313: 179–186.

73. Bujdoso R, Landgraf M, Jackson WS, Thackray AM: Prion-induced neurotoxicity: Possible role for cell cycle activity and DNA damage response. World journal of virology 2015, 4: 188–197.

74. Romphosri S, Changruenngam S, Chookajorn T, Modchang C: Role of a Concentration Gradient in Malaria Drug Resistance Evolution: A Combined within- and between-Hosts Modelling Approach. Scientific Reports 2020, 10: 6219.

75. Nies AT: The role of membrane transporters in drug delivery to brain tumors. Cancer Lett 2007, 254: 11–29.

76. Nies AT, Schwab M, Keppler D: Interplay of conjugating enzymes with OATP uptake transporters and ABCC/MRP efflux pumps in the elimination of drugs. Expert Opin Drug Metab Toxicol 2008, 4: 545–568.

77. Kerb R, Fux R, Mörike K, Kremsner PG, Gil JP, Gleiter CH, Schwab M: Pharmacogenetics of antimalarial drugs: effect on metabolism and transport. Lancet Infect Dis 2009, 9: 760–774.

78. Zuluaga L, Pabón A, López C, Ochoa A, Blair S: Amodiaquine failure associated with erythrocytic glutathione in Plasmodium falciparum malaria. Malaria Journal 2007, 6: 47.

79. Ginsburg H, Golenser J: Glutathione is involved in the antimalarial action of chloroquine and its modulation affects drug sensitivity of human and murine species of Plasmodium. Redox Rep 2003, 8: 276–279.

80. Daoud H, Rouleau GA: A role for ubiquilin 2 mutations in neurodegeneration. Nature Reviews Neurology 2011, 7: 599–600.

81. Qi S, Wang C, Li C, Wang P, Liu M: Candidate genes investigation for severe nonalcoholic fatty liver disease based on bioinformatics analysis. Medicine 2017, 96: e7743–e7743.

82. Wu M, Fang K, Wang W, Lin W, Guo L, Wang J: Identification of key genes and pathways for Alzheimer’s disease via combined analysis of genome-wide expression profiling in the hippocampus. Biophysics Reports 2019, 5: 98–109.

83. Karbalaei R, Allahyari M, Rezaei-Tavirani M, Asadzadeh-Aghdaei H, Zali MR: Protein-protein interaction analysis of Alzheimers disease and NAFLD based on systems biology methods unhide common ancestor pathways. Gastroenterology and Hepatology from bed to bench 2018, 11: 27.

84. Schwabe RF: Endocannabinoids promote hepatic lipogenesis and steatosis through CB1 receptors. Hepatology 2005, 42: 959–961.

85. Jeong WI, Osei-Hyiaman D, Park O, Liu J, Bátkai S, Mukhopadhyay P, Horiguchi N, Harvey-White J, Marsicano G, Lutz B, et al: Paracrine activation of hepatic CB1 receptors by stellate cell-derived endocannabinoids mediates alcoholic fatty liver. Cell Metab 2008, 7: 227–235.

86. Alferink J, Specht S, Arends H, Schumak B, Schmidt K, Ruland C, Lundt R, Kemter A, Dlugos A, Kuepper JM, et al: Cannabinoid Receptor 2 Modulates Susceptibility to Experimental Cerebral Malaria through a CCL17-dependent Mechanism *. Journal of Biological Chemistry 2016, 291: 19517–19531.

87. Cabral GA, Griffin-Thomas L: Emerging role of the cannabinoid receptor CB2 in immune regulation: therapeutic prospects for neuroinflammation. Expert reviews in molecular medicine 2009, 11: e3–e3.

88. Centonze D, Rossi S, Finazzi-Agrò A, Bernardi G, Maccarrone M: The (endo)cannabinoid system in multiple sclerosis and amyotrophic lateral sclerosis. Int Rev Neurobiol 2007, 82: 171–186.

89. Rossi S, Bernardi G, Centonze D: The endocannabinoid system in the inflammatory and neurodegenerative processes of multiple sclerosis and of amyotrophic lateral sclerosis. Experimental Neurology 2010, 224: 92–102.

90. Fowler CJ, Rojo ML, Rodriguez-Gaztelumendi A: Modulation of the endocannabinoid system: neuroprotection or neurotoxicity? Exp Neurol 2010, 224: 37–47.

91. Surowiec I, Gouveia-Figueira S, Orikiiriza J, Lindquist E, Bonde M, Magambo J, Muhinda C, Bergström S, Normark J, Trygg J: The oxylipin and endocannabidome responses in acute phase Plasmodium falciparum malaria in children. Malaria Journal 2017, 16: 358.

92. Son Y, Cheong Y-K, Kim N-H, Chung H-T, Kang DG, Pae H-O: Mitogen-Activated Protein Kinases and Reactive Oxygen Species: How Can ROS Activate MAPK Pathways? Journal of Signal Transduction 2011, 2011: 792639.

93. Puig B, Gómez-Isla T, Ribe E, Cuadrado M, Torrejón-Escribano B, Dalfo E, Ferrer I: Expression of stress-activated kinases c-Jun N-terminal kinase (SAPK/JNK-P) and p38 kinase (p38-P), and tau hyperphosphorylation in neurites surrounding βA plaques in APP Tg2576 mice. Neuropathology and applied neurobiology 2004, 30: 491–502.

94. Chiarini A, Pra ID, Marconi M, Chakravarthy B, Whitfield JF, Armato U: Calcium-sensing receptor (CaSR) in human brain’s pathophysiology: roles in late-onset Alzheimer’s disease (LOAD). Current Pharmaceutical Biotechnology 2009, 10: 317–326.

95. Hashimoto Y, Tsuji O, Niikura T, Yamagishi Y, Ishizaka M, Kawasumi M, Chiba T, Kanekura K, Yamada M, Tsukamoto E: Involvement of c-Jun N-terminal kinase in amyloid precursor protein-mediated neuronal cell death. Journal of neurochemistry 2003, 84: 864–877.

96. Brownlees J, Yates A, Bajaj N, Davis D, Anderton B, Leigh P, Shaw C, Miller C: Phosphorylation of neurofilament heavy chain side-arms by stress activated protein kinase-1b/Jun N-terminal kinase-3. Journal of Cell Science 2000, 113: 401–407.

97. Ackerley S, Grierson AJ, Banner S, Perkinton MS, Brownlees J, Byers HL, Ward M, Thornhill P, Hussain K, Waby JS: p38α stress-activated protein kinase phosphorylates neurofilaments and is associated with neurofilament pathology in amyotrophic lateral sclerosis. Molecular and Cellular Neuroscience 2004, 26: 354–364.

98. Kim EK, Choi E-J: Pathological roles of MAPK signaling pathways in human diseases. Biochimica et Biophysica Acta (BBA) - Molecular Basis of Disease 2010, 1802: 396–405.

99. Wang J-Z, Liu F: Microtubule-associated protein tau in development, degeneration and protection of neurons. Progress in neurobiology 2008, 85: 148–175.

100. Pérez M, Morán MA, Ferrer I, Ávila J, Gómez-Ramos P: Phosphorylated tau in neuritic plaques of APP sw/Tau vlw transgenic mice and Alzheimer disease. Acta neuropathologica 2008, 116: 409–418.

101. Eckermann K, Mocanu M-M, Khlistunova I, Biernat J, Nissen A, Hofmann A, Schönig K, Bujard H, Haemisch A, Mandelkow E: The β-propensity of Tau determines aggregation and synaptic loss in inducible mouse models of tauopathy. Journal of Biological Chemistry 2007, 282: 31755–31765.

102. Alonso AD, Cohen LS, Corbo C, Morozova V, ElIdrissi A, Phillips G, Kleiman FE: Hyperphosphorylation of Tau Associates With Changes in Its Function Beyond Microtubule Stability. Frontiers in Cellular Neuroscience 2018, 12.

103. Dorovini-Zis K, Schmidt K, Huynh H, Fu W, Whitten RO, Milner D, Kamiza S, Molyneux M, Taylor TE: The neuropathology of fatal cerebral malaria in malawian children. Am J Pathol 2011, 178: 2146–2158.

104. Severe malaria. Trop Med Int Health 2014, 19 Suppl 1: 7–131.

105. Medana IM, Day NP, Hien TT, Mai NT, Bethell D, Phu NH, Farrar J, Esiri MM, White NJ, Turner GD: Axonal injury in cerebral malaria. Am J Pathol 2002, 160: 655–666.

106. Kumar SP, Babu PP: Aberrant Dopamine Receptor Signaling Plays Critical Role in the Impairment of Striatal Neurons in Experimental Cerebral Malaria. Mol Neurobiol 2020, 57: 5069–5083.

107. Aarts E, van Holstein M, Cools R: Striatal Dopamine and the Interface between Motivation and Cognition. Frontiers in Psychology 2011, 2.

108. Provost J-S, Hanganu A, Monchi O: Neuroimaging studies of the striatum in cognition Part I: healthy individuals. Frontiers in Systems Neuroscience 2015, 9.

109. Manda-Handzlik A, Demkow U: The Brain Entangled: The Contribution of Neutrophil Extracellular Traps to the Diseases of the Central Nervous System. Cells 2019, 8.

110. Kojima S, Nagamine Y, Hayano M, Looareesuwan S, Nakanishi K: A potential role of interleukin 18 in severe falciparum malaria. Acta Trop 2004, 89: 279–284.

111. Knackstedt SL, Georgiadou A, Apel F, Abu-Abed U, Moxon CA, Cunnington AJ, Raupach B, Cunningham D, Langhorne J, Krüger R, et al: Neutrophil extracellular traps drive inflammatory pathogenesis in malaria. Sci Immunol 2019, 4.

112. Ponts N, Yang J, Chung D-WD, Prudhomme J, Girke T, Horrocks P, Le Roch KG: Deciphering the Ubiquitin-Mediated Pathway in Apicomplexan Parasites: A Potential Strategy to Interfere with Parasite Virulence. PLOS ONE 2008, 3: e2386.

113. Zhu X, Cho ES, Sha Q, Peng J, Oksov Y, Kam SY, Ho M, Walker RH, Lee S: Giant axon formation in mice lacking Kell, XK, or Kell and XK: animal models of McLeod neuroacanthocytosis syndrome. Am J Pathol 2014, 184: 800–807.

114. Tanaka K, Suzuki T, Hattori N, Mizuno Y: Ubiquitin, proteasome and parkin. Biochimica et Biophysica Acta (BBA) - Molecular Cell Research 2004, 1695: 235–247.

