## Supplementary Material for "Inflammation and Cellular Stress Induced Neurological Sequelae of *Plasmodium falciparum* Malaria"

**Supplementary material for the article “Inflammation and Cellular Stress Induced Neurological Sequelae of *Plasmodium falciparum* Malaria”**

Akua A. Karikari[1,3], Wasco Wruck[2,3] and James Adjaye[2]*

[1]Department of Biomedical Sciences, College of Health and Allied Sciences, University of Cape Coast, Ghana

[2]Institute for Stem Cell Research and Regenerative Medicine, Medical Faculty, Heinrich-Heine University, 40225 Düsseldorf, Germany

[3]Akua A. Karikari and Wasco Wruck contributed equally to this work.

**Supplementary table 1 (external file tableS1.xlsx): Venn diagram analysis of CM, NCM and MM.** (a) subsets of genes in the Venn diagram, (b) GO analysis results from the 77 genes expressed exclusively in CM, (c) GO analysis results from the 289 genes expressed exclusively in NCM, (d) GO analysis results from the 66 genes expressed exclusively in MM, (e) KEGG analysis results from the 77 genes expressed exclusively in CM, (f) KEGG analysis results from the 289 genes expressed exclusively in NCM, (g) KEGG analysis results from the 66 genes expressed exclusively in MM. GO analysis results are sorted by the top GO categories Biological Process (BP), Cellular Component (CC) and Molecular Function (MF) and within these categories by p-value.

**Supplementary table 2 (external file tableS2.xlsx): Differential expression analysis of CM vs. NCM.** (a) genes significantly up-regulated (ratio > 2, $p < 0.05$, $q < 0.25$) and down-regulated (ratio < 0.5, $p < 0.05$, $q < 0.25$), (b) GO analysis results from the genes significantly up-regulated in CM, (c) GO analysis results from the genes significantly down-regulated in CM, (d) KEGG pathway analysis results from the genes significantly up-regulated in CM, (e) KEGG pathway analysis results from the genes significantly down-regulated in CM.

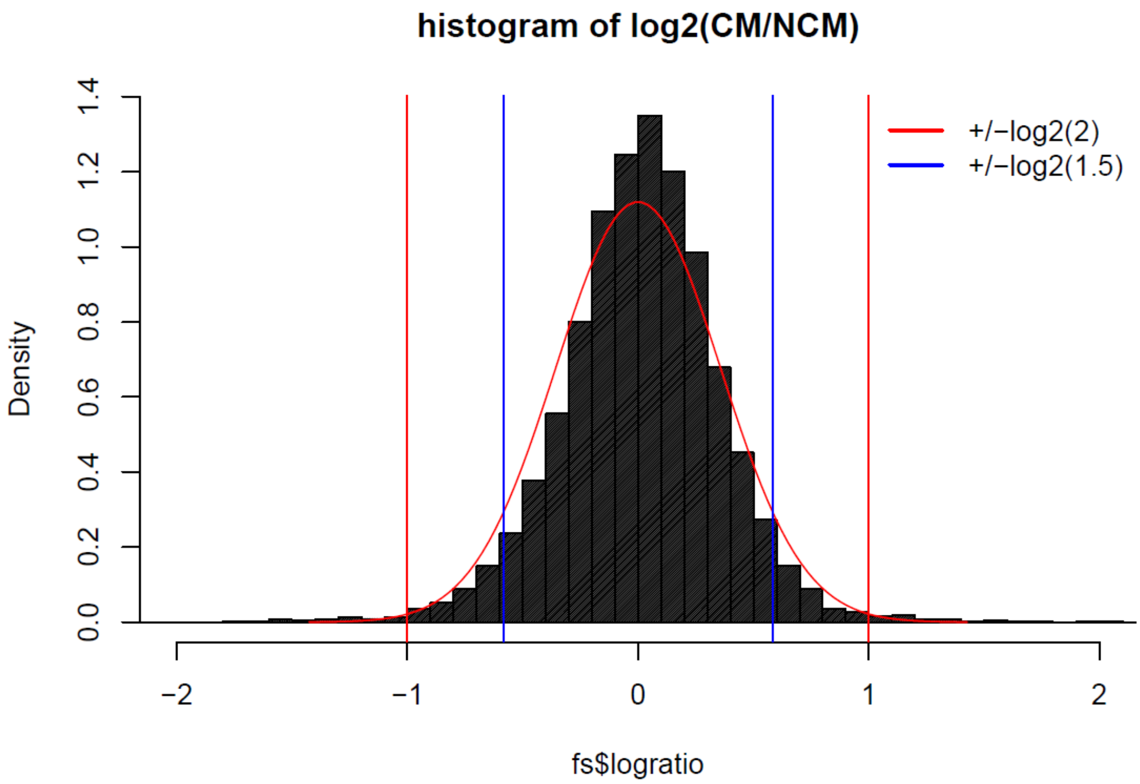


**Supplementary figure 1: Distribution of log2-ratios of CM vs. NCM.** A Gaussian distribution curve is overlaid (mean=0, sd=0.36). Red lines indicate ratio of 2 (0.5) and blue lines ratio of 1.5 (2/3). The area under curve for ratio 1.5 and 2/3 is ~ 90%, the area for ratio 2 and 0.5 is > 99%.

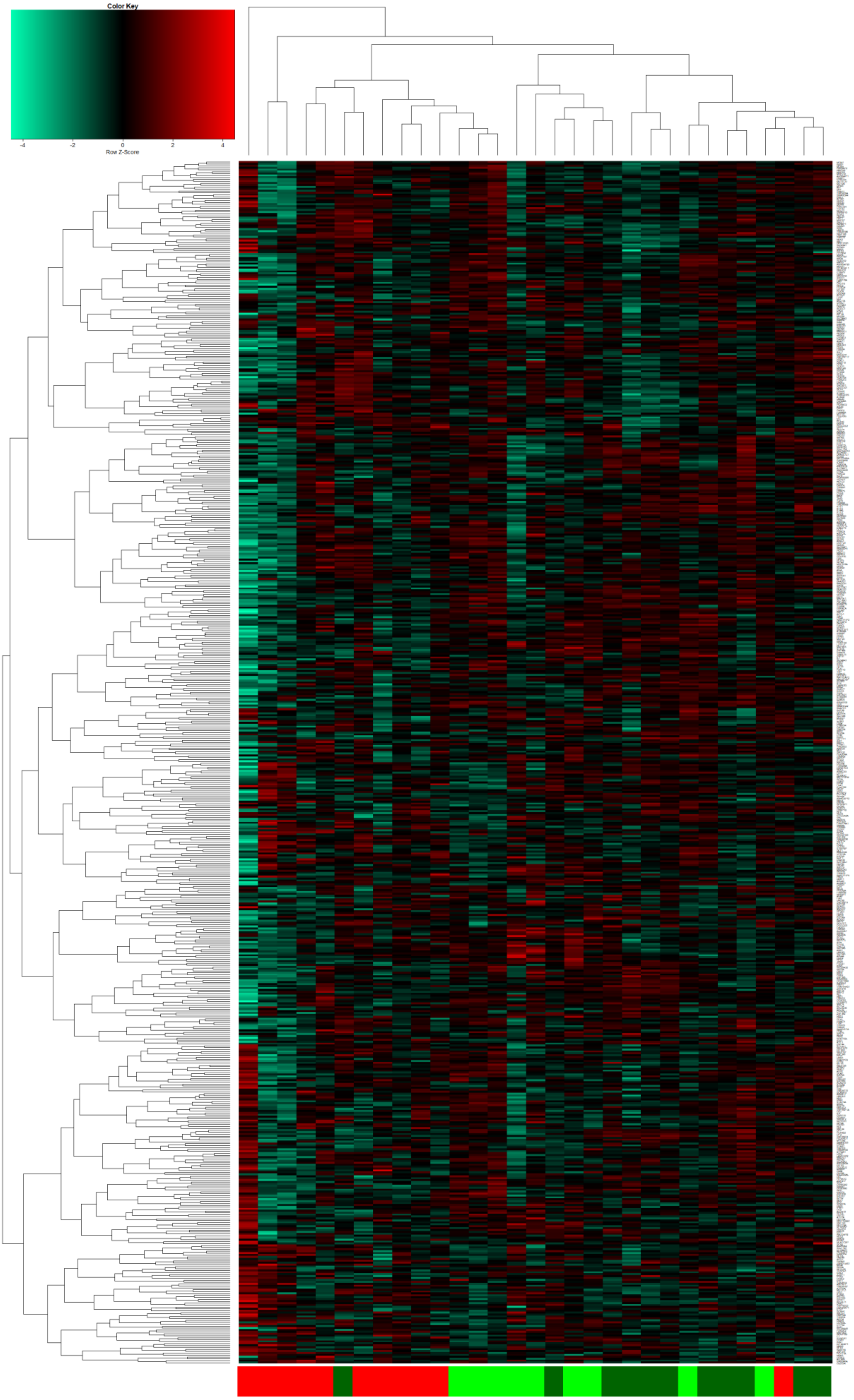
Color Key
-4
-2
0
2
4
Row Z-Score

**Supplementary figure 2: Heatmap and hierarchical cluster analysis of genes expressed exclusively in CM, MM and NCM.** Genes from the exclusive subsets from the Venn diagram depicted in Figure 2c were used to generate a heatmap and perform a hierarchical cluster analysis using Spearman correlation as similarity measure. Color bars indicate: red – CM, green – MM and dark-green - NCM.

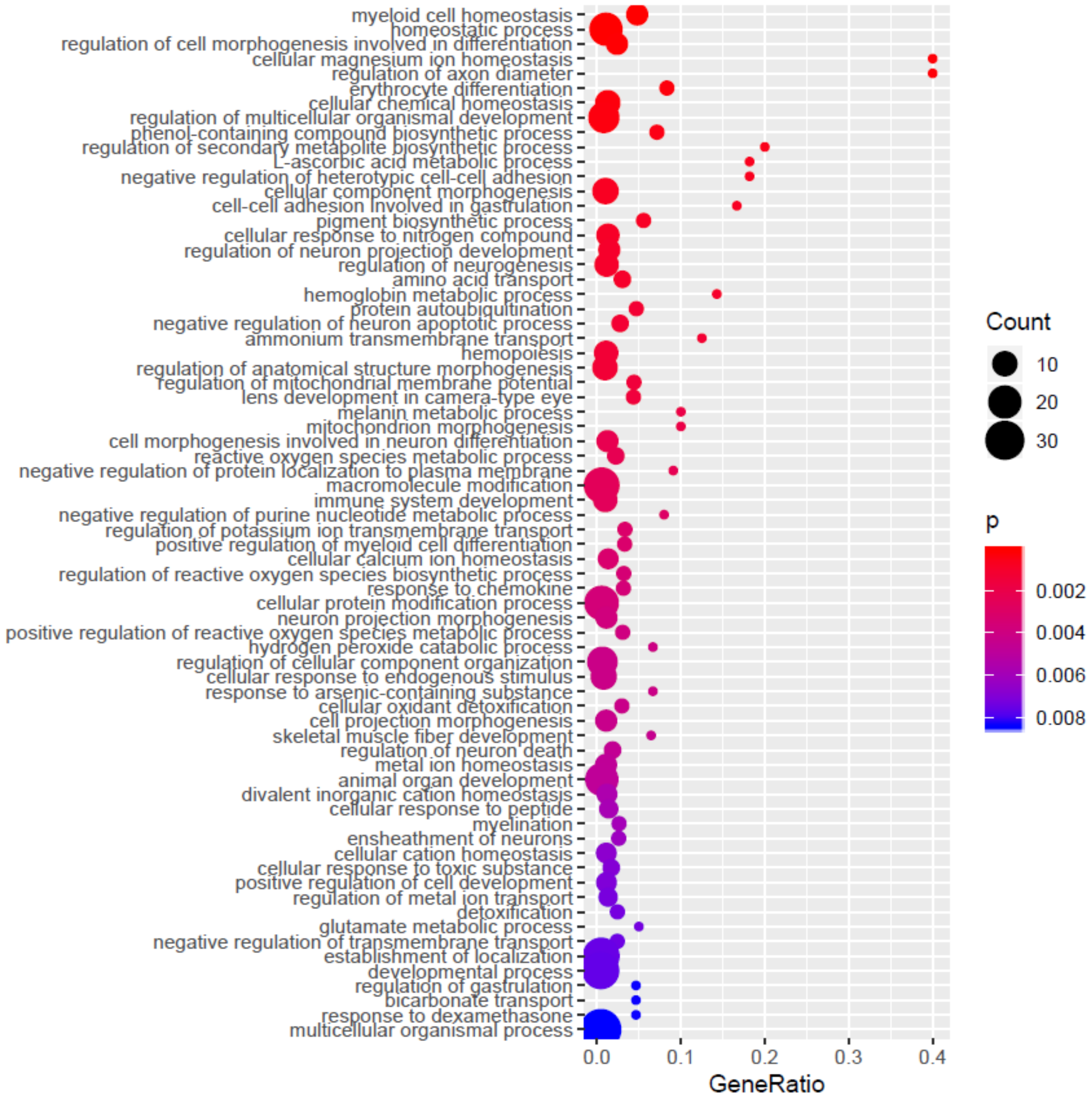


**Supplementary figure 3: 70 most significant GO terms over-represented in genes down-regulated between CM and NCM.** Expansion of Figure 3b which had been reduced to the 30 most significant terms for better readability. Here, all GO (Biological Process) terms below a threshold of 0.01 are depicted, also including *glutamate metabolic process* (see also suppl. Table 2c).
